# *In vitro* Reconstitution Unveils the Phospho-regulated Ska-Ndc80 Macro-complex

**DOI:** 10.1101/680314

**Authors:** Qian Zhang, Liqiao Hu, Yujue Chen, Wei Tian, Hong Liu

## Abstract

The human Ska (<u>s</u>pindle and <u>k</u>inetochore <u>a</u>ssociated) complex is essential for proper kinetochore-microtubule interactions. Its loading onto the kinetochore complex Ndc80 (Ndc80C) plays a critical role in such function; however, the key determinant that docks Ska onto Ndc80C still remain elusive. Here, we address this question using biochemical reconstitution followed by functional analysis. We identified six Cdk1 sites in Ska3 distributed in three conserved regions. *In vitro*, Cdk1 phosphorylation on the Ska complex enhanced WT, not phospho-deficient 6A, binding to Ndc80C. Strikingly, the phospho-mimetic Ska 6D complex formed a stable macro-complex with Ndc80C, but Ska WT failed to do so, suggesting that Cdk1 phosphorylation is necessary and sufficient for the formation of the Ska-Ndc80 macro-complex. In cells, Ska3 6A completely lost its localization to kinetochores and its functions in chromosome segregation; whereas Ska3 6D partially restored them. Altogether, our findings not only for the first time reveal the key phospho-regulated Ska-Ndc80 macro-complex essential for faithful chromosome segregation, but also pave the way for mechanistic analyses of kinetochore-microtubule interactions.

## INTRODUCTION

Proper kinetochore-microtubule interactions are essential for faithful chromosome segregation and preventing chromosome instability. At early mitosis, the kinetochore is initially captured by the spindle microtubules though the KMN (Knl1, the Mis12, and Ndc80 complexes) network, but the initial kinetochore-microtubule interactions on their own are not sufficient to sustain the subsequent chromosome segregation (Cheerambathur and Desai, 2014; Tanaka et al., 2005; Varma and Salmon, 2012). Other critical factors are needed to finalize the KMN-mediated interactions with microtubules. Among them is the Ska complex (Ska), which comprises three subunits, Ska1, 2 and 3 (Guimaraes and Deluca, 2009). Depletion of any subunit in this complex significantly compromised the kinetochore-microtubule interactions and triggered massive mitotic arrest (Daum et al., 2009; Gaitanos et al., 2009; Hanisch et al., 2006; Ohta et al., 2010; Raaijmakers et al., 2009; Theis et al., 2009; Welburn et al., 2009), suggestive of an essential role of Ska in promoting kinetochore-microtubule interactions. However, how this is achieved is not fully understood. In cells, Ska starts to accumulate at kinetochores in prometaphase and peaks at metaphase (Auckland et al., 2017; Hanisch et al., 2006). The dynamic localization pattern coordinates well with the process of establishing proper kinetochore-microtubule interactions from prophase to metaphase, suggesting that Ska binding to kinetochores could be important for its functions. The subsequent findings further supported this notion by demonstrating that the kinetochore receptor for Ska is the Ndc80 complex (Ndc80C, comprising four subunits Ndc80, Nuf2, Spc24 and Spc25) (Chan et al., 2012; Gaitanos et al., 2009; Raaijmakers et al., 2009; Welburn et al., 2009; Zhang et al., 2012), and the physical interaction between Ska and Ndc80C is crucial for Ska functions (Zhang et al., 2012; Zhang et al., 2017). Further studies demonstrated that the Ndc80 N-terminal tail, the calponin-homology domain as well as the internal loop play important roles in recruiting Ska to kinetochores (Cheerambathur et al., 2017; Helgeson et al., 2018; Janczyk et al., 2017; Zhang et al., 2017), suggesting that multiple pathways might exist to install Ska to kinetochores redundantly or collaboratively. However, the key determinant that docks the Ska complex onto kinetochores still remains elusive.

The findings from extensive studies have suggested several potential mechanisms to explain how the Ska complex promotes kinetochore-microtubule interactions. Firstly, loading of Ska to the Ndc80C can bring PP1 to the kinetochore-microtubule binding interface as the Ska1 C-terminus directly binds PP1 that can promote kinetochore-microtubule interactions (Sivakumar et al., 2016). Secondly, Ska binding to Ndc80C may strengthen the kinetochore-microtubule interactions as the C-termini of both Ska1 and Ska3 bear microtubule-binding activities (Abad et al., 2014; Abad et al., 2016; Boeszoermenyi et al., 2014; Jeyaprakash et al., 2012; Thomas et al., 2016). Thirdly, various *in vitro* studies demonstrated that Ndc80C alone or with the help of Ska is able to track with disassembling microtubule tips (Alushin et al., 2010; Monda et al., 2017; Schmidt et al., 2012; Umbreit et al., 2012). All these interesting observations have spotlighted the importance of the physical interaction between Ska and Ndc80C in establishing and/or maintaining proper kinetochore-microtubule interactions, but how these two complexes bind to each other are still not quite understood; and whether they could form a stable complex in vitro is unknown. By analyzing Cdk1 phosphorylation on Ska3, we identified six critical sites in determining the Ska-Ndc80 interaction. *in vitro* reconstitution experiments suggest that Cdk1 phosphorylation on these six sites is necessary and sufficient for the formation of a stable Ska-Ndc80C macro-complex. Further cellular analyses indicate that Cdk1 phosphorylation on these six sites is decisive for Ska localization to kinetochores and Ska functions in chromosome segregation. Thus, our findings for the first time reveal the key phospho-regulated Ska-Ndc80 macro-complex that is essential for proper kinetochore-microtubule interactions and chromosome segregation.

## RESULTS

### The Ska3 C-terminal region is phosphorylated by Cdk1 at multiple sites

In order to understand how the Ska complex interacts with Ndc80C, we attempted to *in vitro* reconstitute the Ska-Ndc80 macro-complex. As Cdk1 phosphorylation promotes Ska3 C-terminal fragments binding to Ndc80C *in vitro* (Zhang et al., 2017), we reasoned that Cdk1 phosphorylation might be the key factor to successfully reconstitute the Ska-Ndc80 macro-complex. Therefore, we in-depth analyzed the fourteen Cdk1 sites located within the Ska3 C-terminal region. We firstly analyzed the conservation of these Cdk1 sites across species, including human (*hs*), mouse (*mm*), chicken (*gg*) and *xenopus laevis* (*xl*) (**Figure S1A**). Six sites, T203, T217, S283, T291, T358, and T360 were found to distribute within three conserved regions. Among them, T203, S283, T291, and T358, are well conserved across the tested species; and the other two sites T217 and T360 are less conserved (**Figures 1A** and **S1A**). Secondly, we performed mass-spectrometric analyses on immunoprecipitated Myc-Ska3 isolated from nocodazole-arrested HeLa Tet-On cells. Among the above six sites, five were identified being phosphorylated in mitotic cells (**Figure 1B)**. T203, although it was not retrieved from our mass-spectrometric analyses, was also included for further analysis because of its high conservation. Finally, all the six sites were able to be phosphorylated by Cdk1 *in vitro* (**Figure S1B**). Thus, these six sites, T203, T217, S283, T291, T358, and T360 were subject to the subsequent analysis in this study. We therefore constructed various mutants containing different sets of mutations on these sites (**Figure 1C)**. As Ska3 phosphorylation significantly reduced its gel mobility on SDS-PAGE (Zhang et al., 2017), we then examined how mutations of these sites affected its migration behavior. Nocodazole-arrested HeLa Tet-On cells were transfected with GFP-Ska3 WT, 2A, 4A-1/2/3, 6A, and 6D, and cell lysates were analyzed with SDS-PAGE followed by Western Blots. Consistent with the previous findings, the slower-migration species of Ska3 WT appeared on SDS-PAGE (**Figure 1D)** (Zhang et al., 2017). Double mutations to alanine (2A) accelerated the gel mobility of the slower species, and quadruple mutations to alanine (4A-1/2) that included Thr358 and Thr360 further accelerated the gel mobility. Strikingly, mutations of all the six sites to non-phosphorylatable alanine (6A) almost completely abolished the slower-migration species; conversely, mutations of them to phospho-mimetic aspartic acid (6D) largely restored them, suggesting that these six sites are the major phosphorylation sites contributing to slower migration in cells. Surprisingly, another quadruple-mutation mutant (4A-3) that excluded T358 and T360 exhibited similar gel mobility to WT, suggesting that these two sites are the major sites contributing to slower migration. Taken together, Ska3 is phosphorylated at multiple sites during mitosis.

**Figure 1.**
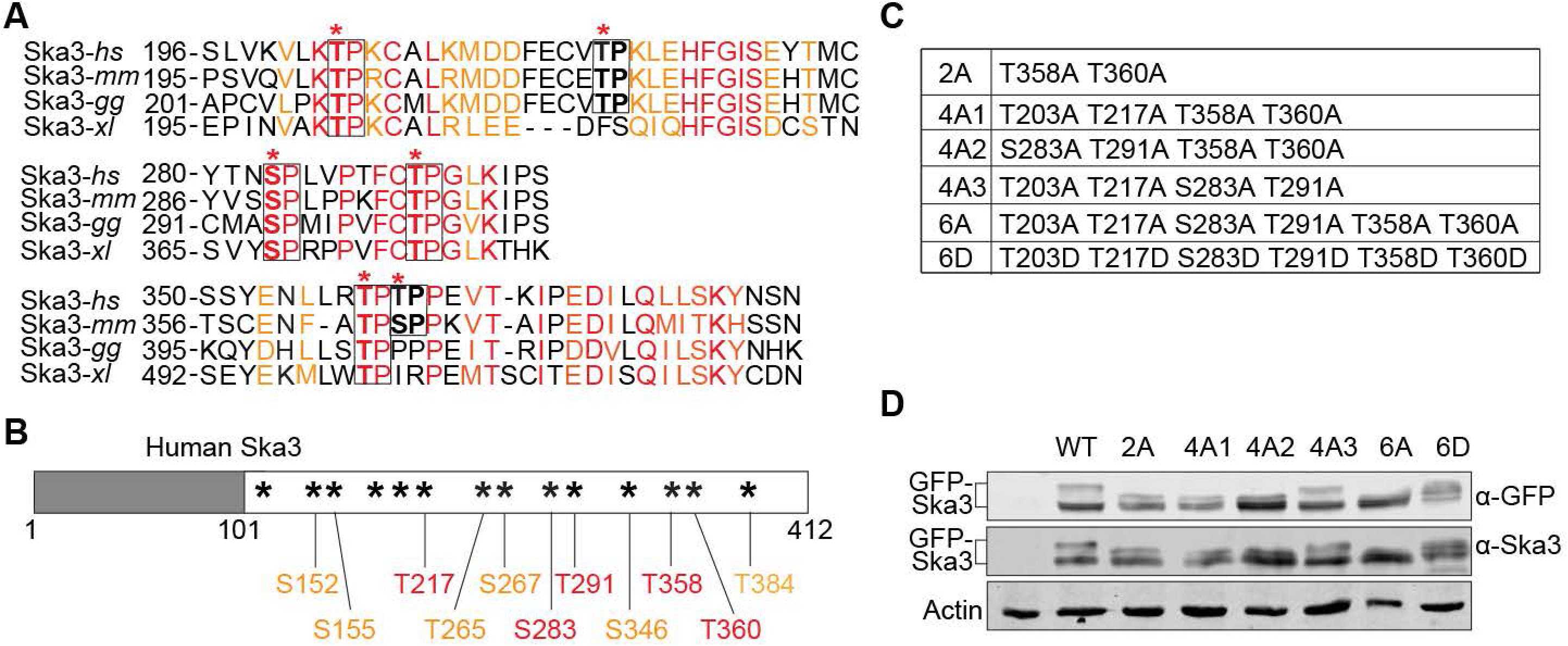
The Ska3 C-terminal region is highly phosphorylated by Cdk1 during mitosis. **A.** Analysis on the conservation of Cdk1 sites in Ska3. Six Cdk1 sites (*) in human Ska3 are distributed in three conserved domains identified from a sequence alignment across four species, human (*hs*), mouse (*mm*), chicken (*gg*), and *xenopus laevis* (*xl*) (**Figure S1**). Among the six sites, four are highly conserved across the tested species. The other two are conserved at least among human and mouse. In the conserved regions, identical amino-acids were marked in red and similar amino-acids in orange. **B.** Phosphorylated Cdk1 sites identified by two mass-spectrometric analyses on immunoprecipitated Myc-Ska3 isolated from nocodazole-arrested HeLa Tet-On cells. The conserved sites were marked in red and the others were marked in orange. Totally eleven Cdk1 sites were identified being phosphorylated. **C.** Summary of Ska3 mutants applied in this study. **D.** Lysates of nocodazole-arrested HeLa Tet-On cells expressing GFP-Ska3 WT, 2A, 4A-1/2/3, 6A, or 6D were resolved with SDS-PAGE and blotted with the indicated antibody.

### Cdk1 phosphorylation on Ska3 promotes the Ska complex binding to Ndc80C in vitro

We next tested how phosphorylation on these six sites affected the interaction between the Ska and Ndc80 complexes. To achieve it, we reconstructed and purified the full-length Ska complex and the GST-Nuf2-Ndc80 complex (**Figure S2**), and performed *in vitro* Cdk1 phosphorylation followed by GST-pull down assays. As shown in **Figure 2A**, an interaction was detected between the Ska complex and GST-Nuf2-Ndc80 even without Cdk1 treatment. This interaction is phospho-independent, which was also suggested in an *in vitro* cross-linking experiment (Helgeson et al., 2018), and its nature is not quite understood. As Cdk1 treatment significantly enhanced the interaction between Ska WT and GST-Nuf2-Ndc80. Treatment of RO3306, a potent Cdk1 inhibitor, completely abolished the enhanced binding between these two complexes, suggesting that the enhanced binding is Cdk1 phosphorylation-dependent. As a comparison, we also examined the interaction between the Ska 6A and GST-Nuf2-Ndc80 complexes under the same conditions. Again, a phospho-independent interaction was detected as well between these two complexes. Either Cdk1 treatment or Cdk1 plus RO3306 treatment did not alter the interaction between Ska 6A and GST-Nfu2-Ndc80 at all. The results from Western Blots using our home-made phospho-specific antibody against T358 and T360 showed that Cdk1 kinase was effective in our experiments (Zhang et al., 2017). Taken the above results together, we concluded that Cdk1 phosphorylation on the six sites in Ska3 promotes the Ska complex binding to Ndc80C.

**Figure 2.**
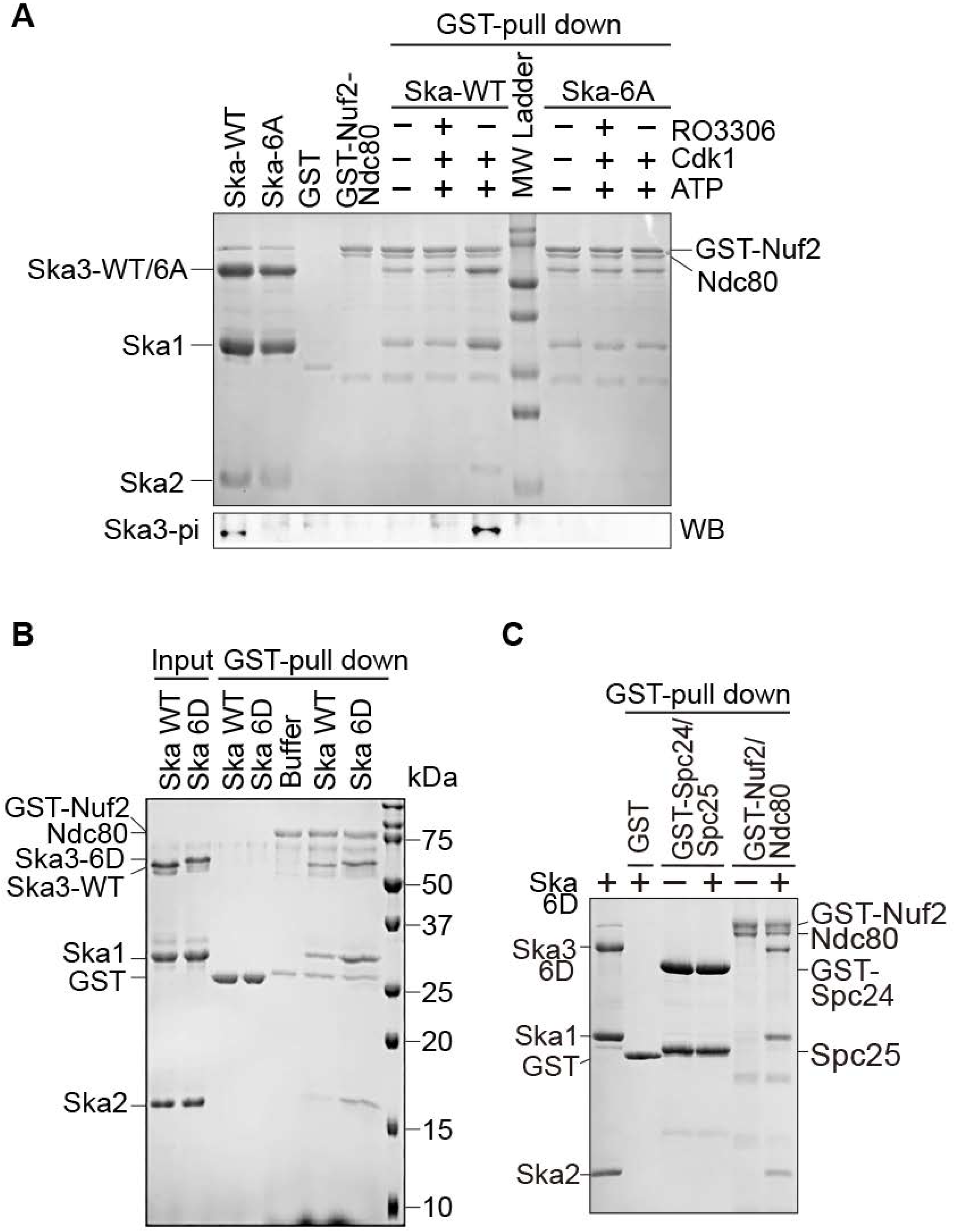
Cdk1 phosphorylation on the six sites in Ska3 promotes the Ska complex binding to Ndc80C. **A.** Cdk1 phosphorylation promotes Ska binding to Ndc80C. The Ska1-Ska2-Ska3 WT or 6A complex treated with Cdk1/Cyclin B1 with or without RO3306 were mixed with the GST-Nuf2-Ndc80 complex and GST beads. The bead-bound proteins were resolved with SDS-PAGE and then stained with Coomassie Blue or blotted with the indicated antibody. **B.** The Ska 6D complex binds to Ndc80C stronger than the WT complex. The Ska1-Ska2-Ska3 WT or 6D complex was mixed with the GST-Nuf2-Ndc80 complex and GST beads. The bead-bound proteins were resolved with SDS-PAGE and then stained with Coomassie Blue. **C.** Spc24 and Spc25 are dispensable for Ska binding. The Ska1-Ska2-Ska3 6D complex was mixed with the GST-Nuf2-Ndc80 or GST-Spc24-Spc25 complex and GST beads. The bead-bound proteins were resolved with SDS-PAGE and then stained with Coomassie Blue.

We then were wondering if the Ska complex containing this phospho-mimetic Ska3 mutant could as well exhibit enhanced binding to Ndc80C. To test this, we examined the interaction between the Ska 6D and GST-Nuf2-Ndc80 complexes using GST-pull down assays. Consistent with our previous results, a phospho-independent interaction was again detected between the Ska WT complex and GST-Nuf2-Ndc80 (**Figure 2B**). Interestingly, an enhanced interaction between the Ska 6D and GST-Nuf2-Ndc80 complexes was also observed, further confirming the phospho-dependent interaction between Ska and Ndc80C. As Ndc80C contains the other two subunits Spc24 and Spc25, we then tested if these two subunits could also interact with the Ska 6D complex. We found that the GST-Spc24-Spc25 complex did not bind the Ska 6D complex at all (**Figure 2C**), suggesting that the subunits Ndc80 and Nuf2 in Ndc80C are mainly responsible for the interaction with the Ska complex.

### The Ska 6D complex forms a stable macro-complex with the Ndc80-Nuf2 complex in vitro

Although we have shown that Cdk1 phosphorylation significantly enhanced the interaction between the Ska and Ndc80 complexes, it is tempting to know whether they could form a stable macro-complex *in vitro*, and if so, could Cdk1 phosphorylation be the decisive factor for the formation of this macro-complex? To overcome the potential heterogeneity of Ska3 phosphorylation by Cdk1, we utilized the phospho-mimetic Ska3 6D mutant as a substitute for phosphorylated Ska3. Because Spc24 and Spc25 are dispensable for Ska binding (**Figure 2C**), only Nuf2-Ndc80 complexes were applied in our *in vitro* reconstitution experiments. We found that the Nuf2-Ndc80 complex was eluted at a high molecular weight, which was in the middle of 669 and 440 kDa (**Figure S2A**). Both Ska WT and 6D complexes were also eluted at high molecular weights, slightly more than 440 kDa (**Figures S2B** and **S2C**). The mixture of the Ska 6D and Nuf2-Ndc80 complexes was eluted as a major peak appearing at a molecular weight slightly less than 440 kDa ((**Figure 3A**). Strikingly, SDS-PAGE analysis on the major peak demonstrated that the Ska 6D and Nuf2-Ndc80 complexes were perfectly co-eluted, revealing the formation of a stable macro-complex. In contrast, the mixture of the Ska WT and Nuf2-Ndc80 complexes was eluted as two major peaks appearing at the molecular weights that were slightly higher and lower than 440 kDa (**Figure 3B**). SDS-PAGE analysis on these two major peaks demonstrated that the Ska WT and Nuf2-Ndc80 complexes were not co-eluted well with each other. All these interesting observations strongly suggest that Cdk1 phosphorylation is the decisive factor for the formation of the stable Ska-Ndc80 macro-complex *in vitro*. Thus, to our knowledge, it is for the first time to successfully reconstitute this macro-complex *in vitro*. In addition, the Ska 6D complex was not co-eluted with the Ndc80 Bosai that lacks the Ndc80 internal loop and the flanking regions (**Figures 3C** **and** **S2D**), although it did so with the full-length Nuf2-Ndc80 complex, further confirming the previous findings that the Ndc80 internal loop together with the flanking regions contains main binding sties for the Ska complex (Zhang et al., 2012; Zhang et al., 2017). Interestingly, the Ndc80 loop is well conserved across species and also responsible for recruiting other important factors (Hsu and Toda, 2011; Maure et al., 2011; Tang et al., 2013; Varma et al., 2012).

**Figure 3.**
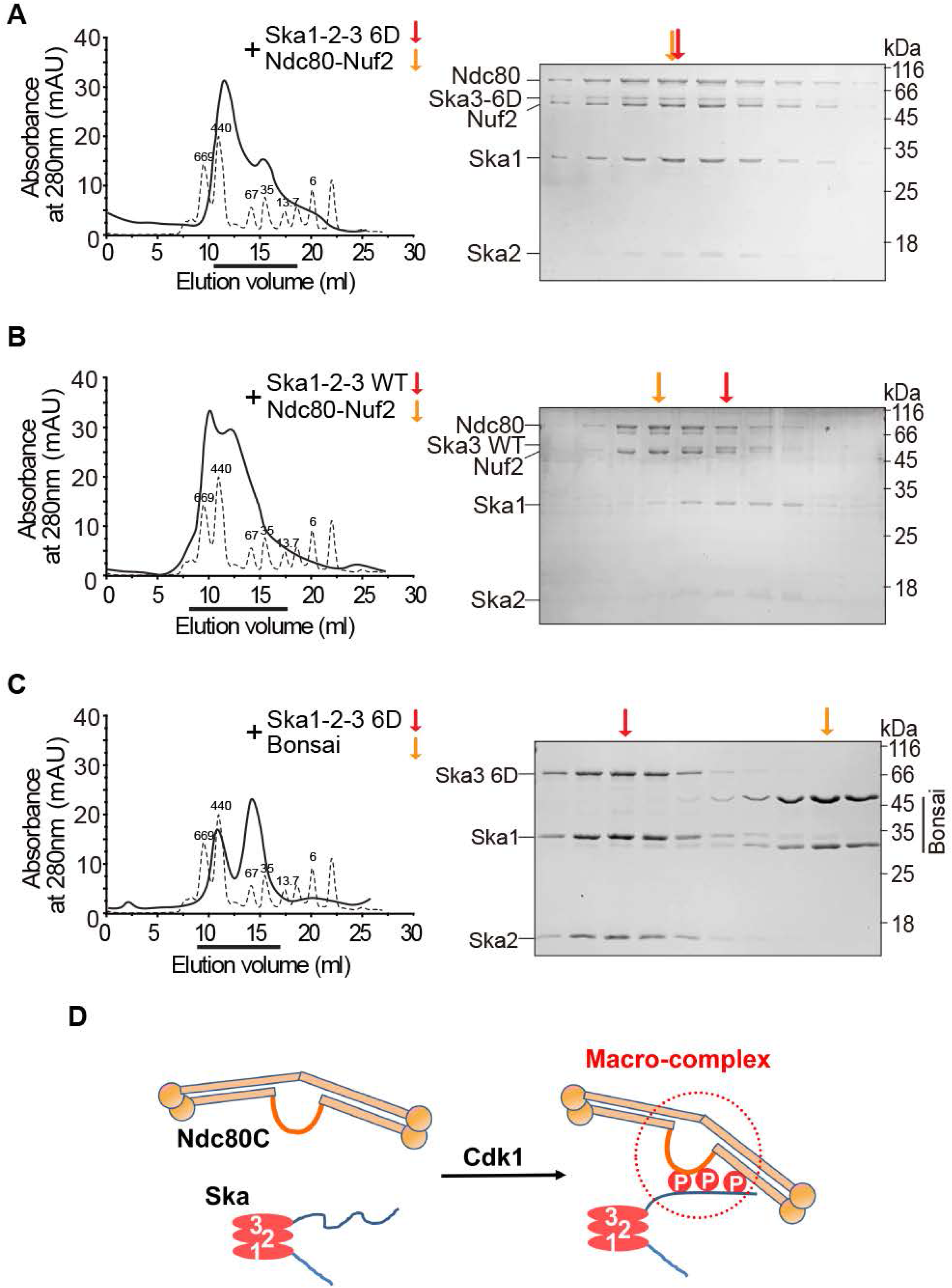
The Ska 6D, not WT complex forms a stable macro-complex with Ndc80C, not Ndc80 Bonsai. **A**, **B** and **C.** Size-exclusion chromatographic analysis on the mixer of the Ska1-Ska2-Ska3 6D and Nuf2-Ndc80 complexes (**A**), or the one of the Ska1-Ska2-Ska3 WT and Nuf2-Ndc80 complexes (**B**), or the one of the Ska1-Ska2-Ska3 6D and the Ndc80 Bosai (**C**). The thicker lines marked the eluted fractions that were analyzed by SDS-PAGE. Arrows indicated the peaks of the analyzed complexes. **D.** Working model. Cdk1 phosphorylation on the Ska3 C-terminal region promotes the formation of a stable Ska-Ndc80 macro-complex. The outlined region with the dotted circle showed the binding domains between the Ska and Ndc80 complexes.

### Cdk1 phosphorylation on the six sites in Ska3 plays a decisive role in Ska3 localization to kinetochores

We have shown that Cdk1 phosphorylation on the six sites is sufficient for promoting the formation of the Ska-Ndc80 macro-complex. We next sought to determine how it affected Ska localization to kinetochores. As Ska localization to kinetochores is dynamically regulated, we then examined Ska3 kinetochore localization in unperturbed, nocodazole-treated and MG132-treated mitotic cells. Thymidine-arrested HeLa Tet-On cells expressing GFP-Ska3 WT, 2A, 4A-1/2/3, 6A, and 6D, were released to fresh medium and mitotic cells were collected for immunostaining. As shown in **Figures 4A** **and** **4B**, the fluorescence signals of GFP-Ska3 WT were detected at kinetochores in prometaphase and metaphase cells. The signals of GFP-Ska3 2A were greatly reduced, consistent with our previous results (Zhang et al., 2017). The ones of GFP-Ska3 4A-1 and 4A-2 that included the mutations of T358 and T360 were also significantly reduced compared to WT. Strikingly, GFP-Ska3 6A with all the six sites mutated to alanine completely lost its kinetochore signals. Conversely, the signals of the phospho-mimetic GFP-Ska3 6D were readily detected at kinetochores albeit weaker than the ones of WT. All these observations support the notion that Cdk1 phosphorylation determines Ska localization to kinetochores. The observed localization defects for these mutants are unlikely due to distinct protein expression levels as they were all comparable to WT (**Figure 1D**). Interestingly, GFP-Ska3 4A-3 that excluded the mutations of T358 and T360 was still localized to kinetochores as robustly as WT (**Figures 4A** and **4B**), suggesting that our previously identified Thr358 and Thr360 are the major sites contributing to Ska kinetochore localization (Zhang et al., 2017). Of note, Similar localization patterns for these mutants were also observed in MG132-arrested metaphase cells (**Figures S3A** and **S3B**). In addition, live-cell imaging also demonstrated that GFP-Ska3 WT, not 6A, was localized to kinetochores although both of them appeared on spindle microtubules (**Figure 4E**). Thus, Cdk1 phosphorylation on Ska3 is the decisive factor for Ska localization to kinetochores during mitosis.

**Figure 4.**
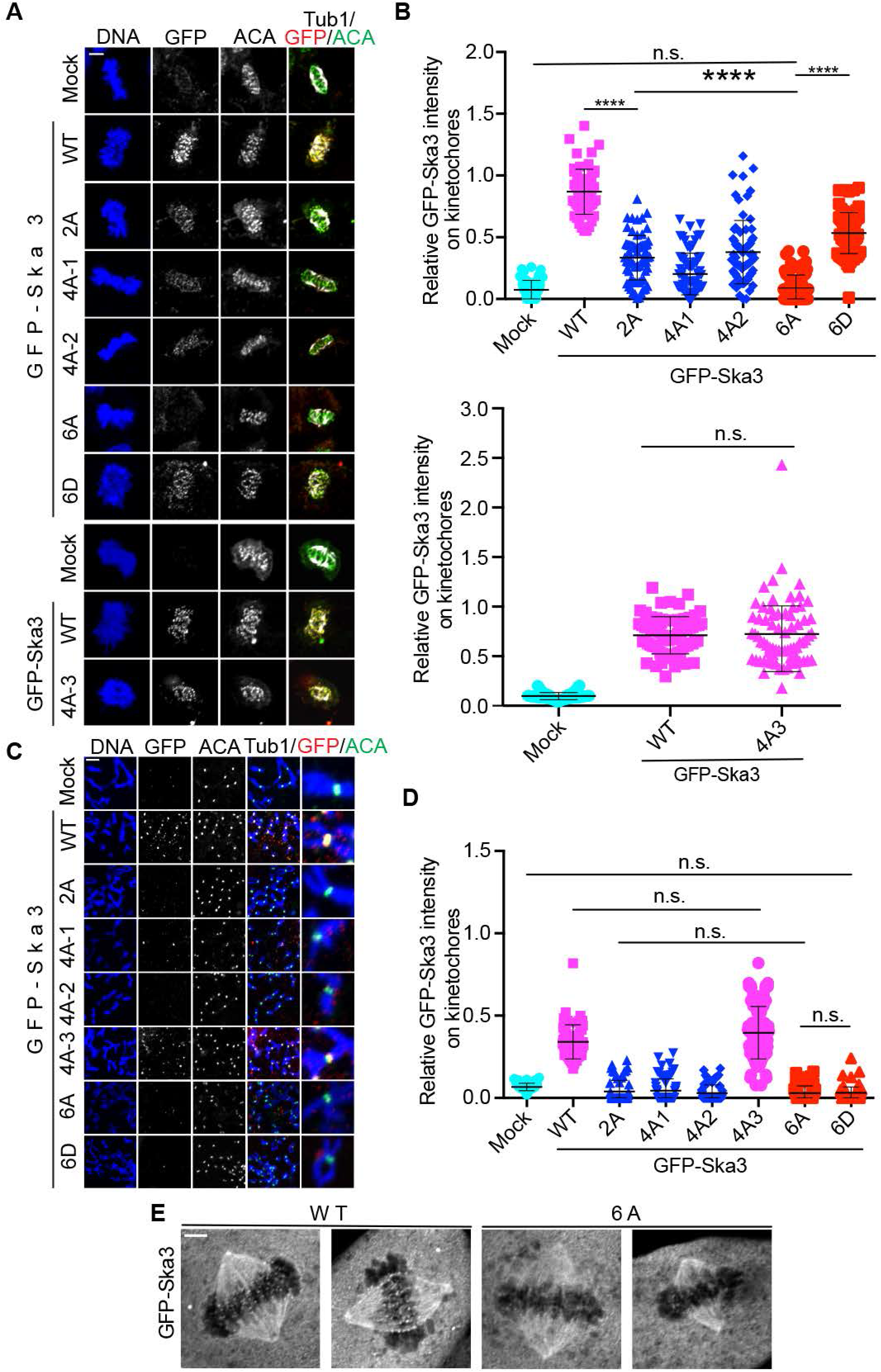
Mutations of all the six Cdk1 sites to alanine (6A) abolish Ska3 localization to kinetochores. **A.** Thymidine-arrested HeLa Tet-On cells expressing GFP-Ska3 WT, 2A, 4A-1/2/3, 6A, or 6D were released into fresh medium. Mitotic cells were collected for immunostaining using the indicated antibodies. Representative images were shown here. Scale bars, 5 μm. **B.** Quantification of GFP-Ska3 intensity on kinetochores in (**A**). Detailed description about quantification was recorded in the section of Methods. At least 50 kinetochores (6 kinetochores per cell) were analyzed for each condition. Average and standard deviation were shown in lines. P<0.0001 (****). n.s. denotes no significance. **C.** Cells were treated similarly to the ones in (A) expect for 2 hr nocodazole-treatment before harvest. Mitotic cells collected were subject to chromosome spread and immunostaining. Scale bars, 5 μm. **D.** Quantification of GFP-Ska3 intensity on kinetochores in (**C**). Detailed description about quantification was recorded in the section of Methods. At least 50 kinetochores (6 kinetochores per cell) were analyzed for each condition. Average and standard deviation were shown in lines. n.s. denotes no significance. **E.** Live-cell imaging of HeLa Tet-On cells expressing GFP-Ska3 WT or 6A treated with MG132. Scale bars, 5 μm.

We next examined the kinetochore localization of these mutants in nocodazole-arrested prometaphase cells. GFP-Ska3 WT, 2A, 4A-1/2/3, 6A, and 6D, were expressed in nocodazole-treated HeLa Tet-On cells and their kinetochore localization was examined. In the presence of nocodazole, GFP-Ska3 WT was detected at kinetochores. Interestingly, all the tested mutants except for 4A-3 completely lost the kinetochore signals (**Figures 4C** and **4D**). Considering that all the mutants except for 4A-3 had more or less defects in kinetochore localization during the unperturbed mitosis (**Figures 4A** and **4B**), the above observations may suggest that the remaining kinetochore binding activities in these mutants might not be sufficient to resist the kinetochore-removing signals imposed by nocodazole treatment. Similar results were also obtained in cells depleted of endogenous Ska3 (**Figures S3C** and **S3D**). Thus, all our results here reveal a decisive role of Cdk1-mediated multisite phosphorylation in Ska localization to kinetochores. The contrasting behavior of 4A-3 further confirmed that our previously identified Thr358 and Thr360 are major sites contributing to Ska localization to kinetochores (Zhang et al., 2017).

### Cdk1 phosphorylation on the six sites in Ska3 is required for proper chromosome segregation

As the Ska complex is required for chromosome segregation, we next determined how the Cdk1 phosphorylation on Ska3 affected chromosome segregation. GFP-Ska3 WT, 6A and 6D were expressed in Ska3-depleted HeLa Tet-On cells stably expressing mCherry-H2B, and then chromosome dynamics was monitored by time-lapse microscopy. As shown in **Figures 5A** and **5B**, Mock cells spent an average of 50 min from nuclear envelop breakdown (NEB) to anaphase onset. Depletion of Ska3 gave rise to two major groups of cells — the one with prolonged metaphase followed by anaphase onset and the other with prolonged metaphase followed by cohesion fatigue and cell death. We quantified mitotic duration (from NEB to anaphase onset or from NEB throughout cohesion fatigue to cell death) that these two groups of cells spent and found an average of 337 min. Expression of GFP-Ska3 WT largely rescued the defects caused by Ska3 depletion with an average mitotic duration of 59 min, which exhibited no significant difference from mock cells. Strikingly, expression of GFP-Ska3 6A completely failed to rescue the defects with an average mitotic duration of 261 min, which was not statistically different from the one for Ska3 depletion. As Ska3 6A completely lost its kinetochore localization (**Figure 4**), the above results strongly support the notion that Ska localization to kinetochore is crucial for its functions in chromosome segregation. These results also reveal a decisive role of Cdk1 phosphorylation in activating Ska functions. In addition, we also found that expression of phospho-mimetic mutant GFP-Ska3 6D partially restored Ska functions in chromosome segregation with an average mitotic duration of 135 min, which was statistically different from the one for 6A. The reason that Ska3 6D only partially rescued the defects will be discussed in the section of Discussion.

**Figure 5.**
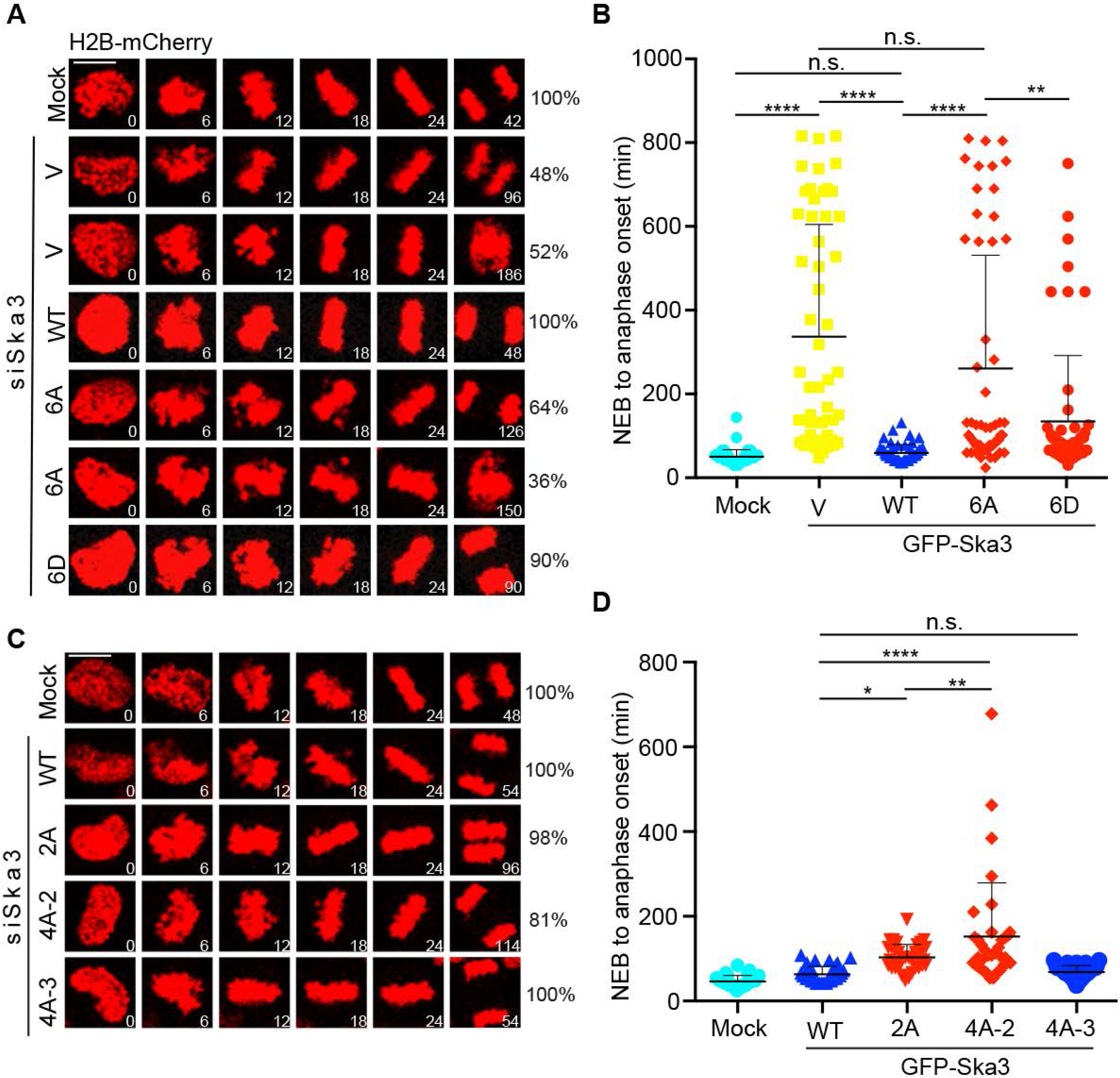
Mutations of all the six Cdk1 sites to alanine (6A) disable Ska functions in chromosome segregation. **A** and **C.** HeLa Tet-On cells stably expressing H2B-mCherry were co-transfected with siSka3 and vectors (V) or plasmids containing GFP-Ska3 WT, 2A (**C**), 4A-2 (**C**), 4A-3 (**C**), 6A (**A**), or 6D (**A**). Time-lapse microscopic analysis was performed. Scale bars, 5 μm. **B** and **D.** Quantification of the duration from nuclear envelop breakdown (NEB) to anaphase onset in (**A**) and (**C**). At least 54 mitotic cells in (**A**) and 50 mitotic cells in (**C**) were analyzed for each condition. Average and standard deviation were shown in lines. P<0.05 (*), P<0.01 (**), P< 0.0001 (****). n.s. denotes no significance. Scale bars, 5 μm.

We also examined chromosome segregation in siSka3-treated cells expressing GFP-Ska3 2A, 4A-2 and 4A-3 (**Figures 5C** and **5D**). Consistent with our previous findings, expression of GFP-Ska3 2A significantly delayed chromosome segregation with an average mitotic duration of 116 min (Zhang et al., 2017). Cells expressing GFP-Ska3 4A-2 that included the mutations of T358A and T360A were also delayed in chromosome segregation with an average mitotic duration of 155 min. Strikingly, expression of GFP-Ska3 4A-3 that excluded the mutations of T358A and T360A did not delay chromosome segregation with an average mitotic duration of 72 min, which did not show any significant difference from GFP-Ska3 WT (65 min). Again, these results support that Thr358 and Thr360 are the major sites contributing to Ska functions. Of note, our results also suggest that the Ska complex may be dispensable for chromosome congression to metaphase plates from initial attachments (**Figures S4A** and **S4B**), consistent with the previous findings (Auckland et al., 2017; Daum et al., 2009). The significant delay in our experiments was derived from metaphase-to-anaphase transition.

## DISCUSSION

An outstanding question in the field is how proper kinetochore-microtubule interactions are achieved during mitosis. Extensive efforts have spotlighted several key players in the kinetochore-microtubule interface. Among these key players is the Ska complex; however, how it promotes proper kinetochore-microtubule interactions is not quite understood. Our findings here provide strong evidence to support that Cdk1 phosphorylation is required for docking Ska to kinetochores and the phosphorylation-mediated kinetochore binding activates Ska functions in kinetochore-microtubule interactions. Recent studies indicated that the Ndc80 tail and/or the calponin-homology domain facilitate Ska recruitment to kinetochores (Cheerambathur et al., 2017; Janczyk et al., 2017), which appears to be inconsistent with our findings that the Ndc80 internal loop together with the flanking regions is the major Ska binding site (Zhang et al., 2017). This seeming discrepancy might be due to the existence of multiple pathways responsible for installing Ska to kinetochores. Alternatively, considering the decisive roles of Cdk1 phosphorylation and the Ndc80 internal loop in recruiting Ska to kinetochores, we prefer another explanation that the Ndc80 N-terminal tail and/or the calponin-homology domain may facilitate the initial Ska recruitment to the kinetochore-microtubule interface, and then the Ndc80 internal loop might finally dock Ska to kinetochores (**Figure 3D**). In future, it would be tempting to test if this is the case.

Why is multisite phosphorylation needed? The Ska complex starts to accumulate at kinetochores in prophase and peaks at metaphase (Auckland et al., 2017; Hanisch et al., 2006), suggesting that the binding affinity of Ska to kinetochores varies with the mitotic progression. Multisite phosphorylation may provide such regulation with distinct sets of combinations. In support of this idea, we found that the state of Cdk1 phosphorylation on Ska3 is positively correlated with the robustness of Ska3 localization to kinetochores. Although mutations on all the six Cdk1 sites abolished Ska localization to kinetochores and its functions in chromosome segregation, their contributions seem to be different. Mutations (2A and 4A-1/2) that include the two sites T358 and T360 are always associated with the phenotypes of decreased kinetochore localization and prolonged mitotic progression; whereas mutations (4A-3) that exclude T358 and T360 exhibit no detectable defects in kinetochore localization and normal mitotic progression. Thus, phosphorylation on the two sites Thr358 and Thr360 likely lays a foundation for Ska localization to kinetochores; and the subsequent phosphorylation on other sites could further enhance its kinetochore recruitment. In addition to Cdk1 phosphorylation, it was shown that treatment of a microtubule-depolymerizing drug significantly reduced Ska localization to kinetochores (Chan et al., 2012), suggesting that either microtubule attachments to kinetochores facilitate Ska loading to kinetochores or Aurora B negatively regulates Ska localization to kinetochores, as proposed (Chan et al., 2012). Thus, it is likely that various mechanisms in cells may collaborate to ensure the precise Ska loading to kinetochores (Zhang et al., 2018). Of note, although the Ska 6D complex bound to Ndc80C stronger than WT, it exhibited partial defects in kinetochore localization and chromosome segregation, suggesting that Ska3 6D might not be a perfect mimicry to phosphorylated Ska3 in cells. Alternatively, as a recent study showed dynamic phosphorylation-dephosphorylation cycles on Ska3 is important proper Ska functions (Maciejowski et al., 2017), it is possible that similar phosphorylation-dephosphorylation cycles may also be critical to efficiently load the Ska complex to kinetochores (Redli et al., 2016; Sivakumar and Gorbsky, 2017).

Our *in vitro* reconstitution results showed the phospho-mimetic Ska 6D complex was able to form a stable macro-complex with the Ndc80-Nuf2 complex, suggesting that a tight Ska-Ndc80 interaction can be formed in cells albeit its existence might be transient. As the Ndc80 internal loop together with its flanking regions is mainly responsible for Ska binding and the loop confers structural flexibility to Ndc80C (Varma et al., 2012; Wang et al., 2008; Zhang et al., 2012; Zhang et al., 2017), such a tight Ska-Ndc80 interaction might be able to alter the conformation of Ndc80C, thus promoting proper kinetochore-microtube interactions. In this sense, the Ska complex not only functions as a scaffold to bring PP1 or other factors in proximity to the kinetochore-microtubule interface, but also might serve as a structural modifier. In future, it is tempting to test if this could be the case; and if so, how does it affect the behavior of Ndc80C on dynamic microtubules (Ye and Maresca, 2013)? In budding yeast, the functional Ska homolog is the DASH/Dam1 complex, which contains 10 subunits and shares no structural and sequence similarity to the Ska complex. The Dam1 complex was reconstituted *in vitro* and its structure was solved using cryo-EM (Jenni and Harrison, 2018). It was also shown that the Ndc80 complex bridges two Dam1 complex rings (Kim et al., 2017). Although integrating the structure of the Ndc80 complex and published interaction data into that of the Dam1 complex yielded an interesting molecular view of kinetochore-microtubule interactions (Jenni and Harrison, 2018), how these complexes indeed interacted with each other still remains unclear. In future, by solving the structure of the Ska (6D)-Ndc80 macro-complex using cryo-EM, people would gain profound insights into how Cdk1 phosphorylation on Ska3 promotes the Ska-Ndc80 interaction at the atomic level.

In summary, our findings reveal an essential role of Cdk1 phosphorylation in controlling Ska localization and functions during mitosis, and for the first time unveil the key phospho-regulated Ska-Ndc80 macro-complex that is essential for proper kinetochore-microtubule attachments and chromosome segregation (**Figure 3D**).

## AUTHOR CONTRIBUTIONS

Conceptualization, H.L.; Methodology, Q.Z., LQ.H., YJ.C., W.T., and H.L.; Investigation, Q.Z., LQ.H., YJ.C., W.T., and H.L.; Writing – Original Draft, Q.Z., H.L.; Writing – Review & Editing, Q.Z., LQ.H., YJ.C., W.T., and H.L.; Funding Acquisition, W.T. and H.L.; Resources, W.T. and H.L.; Supervision, W.T. and H.L..

## ACKNOWLEDGEMENT

This work was supported by the Tulane startup funds awarded to H.L., and by the National Natural Science Foundation of China (31470763 and 31500629) to awarded to W.T.. We also thank Dr. Iain Cheeseman (MIT) for the plasmids containing Ska1 and Ska2.

The authors declare that they have no conflict of interest.

---

**Figure S1.**
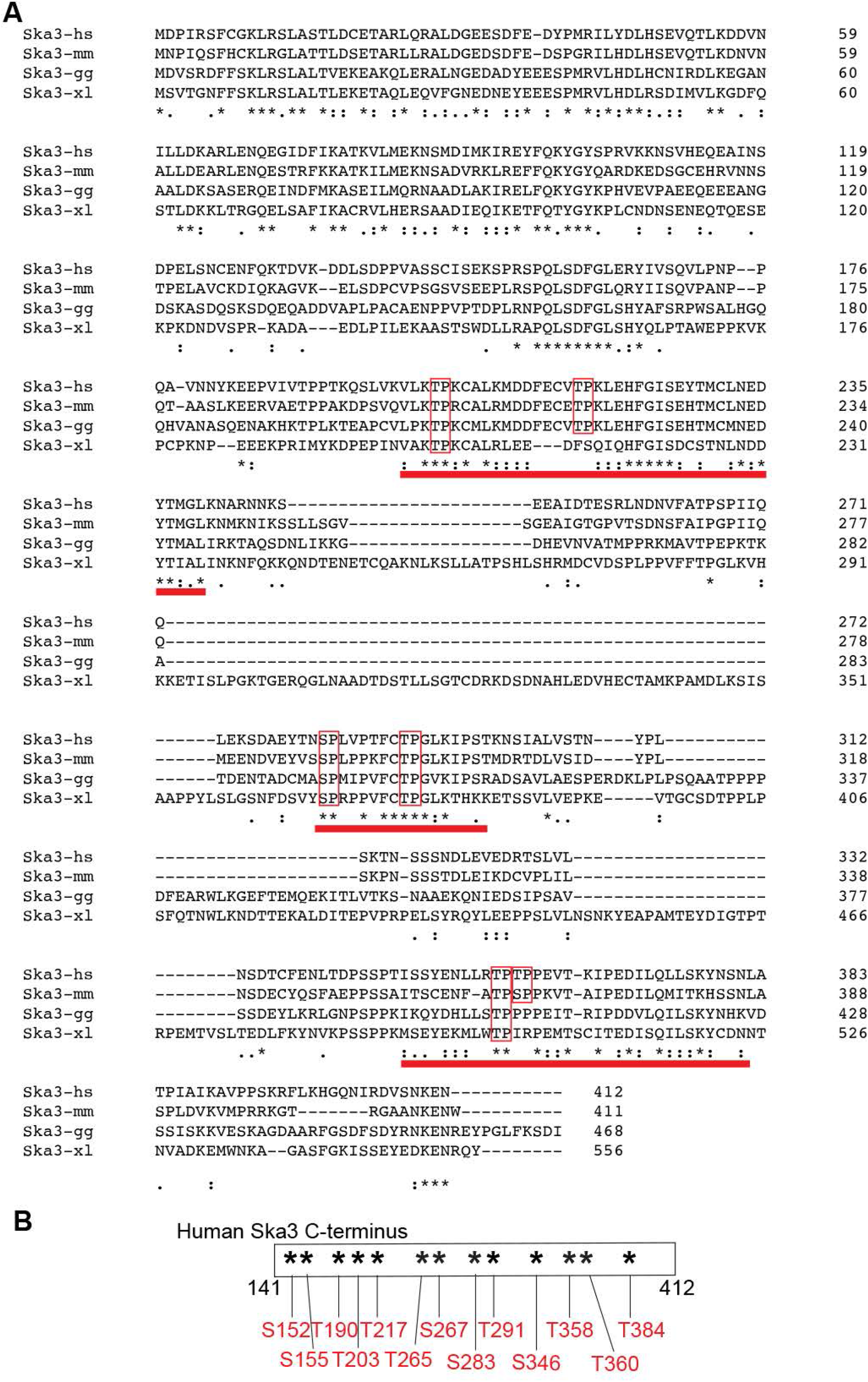
Alignment of Ska3 protein sequences across species. **A.** Ska3 protein sequences among human (*hs*), mouse (*mm*), chicken (*gg*), and *xenopus laevis* (*xl*) were analyzed using Clustal Omega. Identical amino-acid residues were marked with stars and similar amino-acid residues were marked with two dots. Three conserved domains were identified and underscored in red. Conserved Cdk1 sites within these domains were outlined in red. **B.** Phosphorylated Cdk1 sites identified by mass-spectrometric analysis on recombinant Ska3 fragments (141-412) treated with Cdk1/Cyclin B1 *in vitro*.

**Figure S2.**
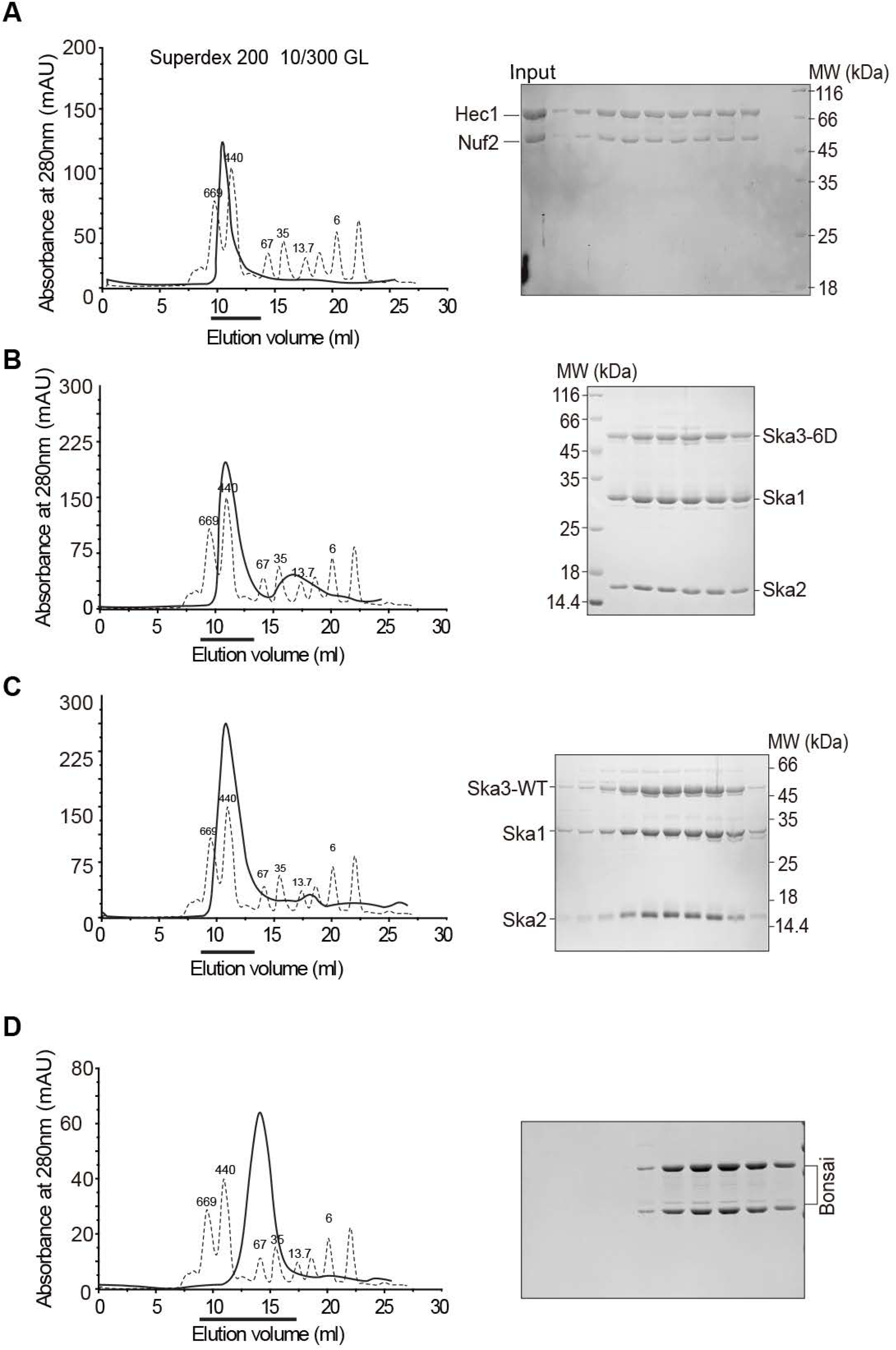
Purification of the Ska and Ndc80 complexes. Size-exclusion chromatographic analysis on purified Nuf2-Ndc80 complexes (**A**), Ska1-Ska2-Ska3 6D complexes (**B**), Ska1-Ska2-Ska3 WT complexes (**C**) and Ndc80 Bosai (**D**). The thicker lines marked the eluted fractions that were analyzed by SDS-PAGE.

**Figure S3.**
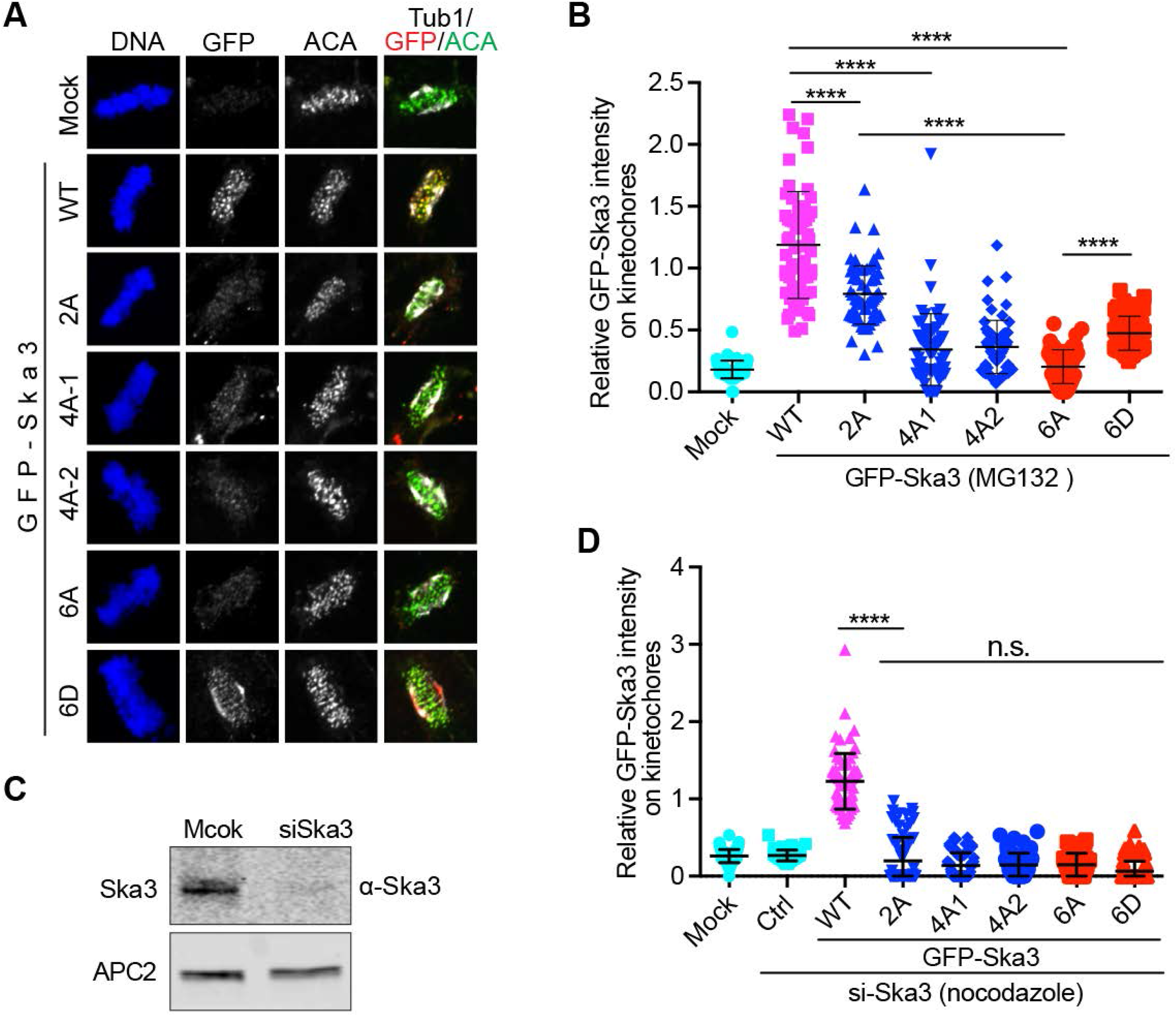
Localization of Ska3 mutants in MG132- and siSka3-treated cells. **A.** Thymidine-arrested HeLa Tet-On cells expressing GFP-Ska3 WT, 2A, 4A-1/2, 6A, or 6D were released into fresh medium. Mitotic cells were harvested after 2-hour MG132 treatment and then subject to immunostaining using the indicated antibodies. Representative images were shown here. **B.** Quantification of GFP-Ska3 intensity on kinetochores in (**B**). Detailed description about quantification was recorded in the section of Methods. Detailed description about quantification was recorded in the section of Methods. At least 50 kinetochores (6 kinetochores per cell) were analyzed for each condition. Average and standard deviation were shown in lines. P<0.0001 (****). n.s. denotes no significance. **C.** Lysates of HeLa Tet-On cells with mock or siSka3 treatment were resolved with SDS-PAGE and blotted with the indicated antibody. APC2 served as loading control. **D.** Quantification of GFP-Ska3 intensity on kinetochores in siSka3-treated HeLa Tet-On cells expressing GFP-Ska3 WT, 2A, 4A-1/2, 6A, or 6D. Detailed description about quantification was recorded in the section of Methods. At least 50 kinetochores (6 kinetochores per cell) were analyzed for each condition. Average and standard deviation were shown in lines. P<0.0001 (****). n.s. denotes no significance.

**Figure S4.**
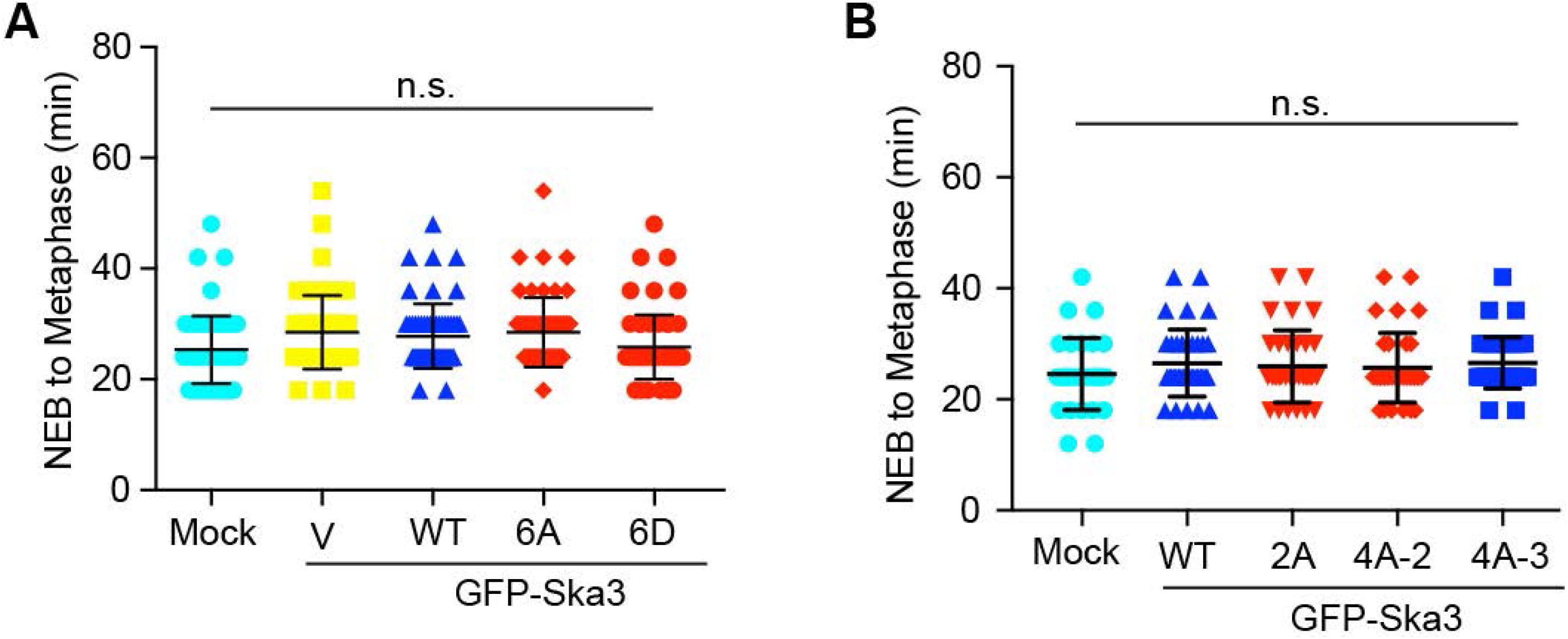
**A** and **B.** Duration from NEB to metaphase for cells in **Figures 5A** and **5C**.

## METHODS AND MATERIALS

### Mammalian cell culture, siRNAs and transfection

HeLa Tet-On cells (Clontech) were cultured in Dulbecco’s modified Eagle’s medium (DMEM, Invitrogen) supplemented with 10% fetal bovine serum and 10mM L-glutamine. To arrest cells at G1/S, cells were incubated in medium containing 2mM thymidine (Sigma) for 16 h. Nocodazole, MG132 and RO3306 were purchased from Sigma Aldrich.

Plasmid transfection was done using the Effectene reagent (Qiagen) according to the manufacturer’s protocols. For H2B-mCherry stable cells, HeLa Tet-On cells were transfected with pIRES vectors encoding H2B-mCherry and selected with 0.4 μg ml^−1^ puromycin (Invitrogen). For Myc-Ska3 stable cells, HeLa Tet-On cells were transfected with pTRE2 vectors encoding RNAi-resistant MYC-Ska3 and selected with 350 μg ml^−1^ hygromycin (Invitrogen). The surviving clones were screened for expression of the desired proteins in the presence of 1μg ml^−1^ doxycycline (Invitrogen). Expression of Myc-Ska3 was also induced with 1μg ml^−1^ doxycycline in the subsequent experiments.

For RNAi experiments, the siRNA oligonucleotides were purchased from Thermo Scientific. HeLa cells were transfected using Lipofectamine RNAiMax (Invitrogen) and analyzed at 32-40 h after transfection. The sequences of the siRNAs used in this study are: Ska3 siRNA, GGAUAUAGUCCACGUGUCA (D-015700-17, MQ-015700-01-0002, Thermo Scientific).

### Antibodies and Immunoblotting and Immunoprecipitation

Antibodies used in this study were listed in the following: Anti-centromere antibody (ACA or CREST-ImmunoVision, HCT-0100), anti-Tubulin (Thermo Scientific, 62204), anti-Actin (Thermo Scientific, MA5-11869) and anti-Ska3 (Bethyl, A304-215A), anti-GFP (Abcam, ab1218; Aves, GFP-1010). Anti-pSka3 were made in-house as described previously (Zhang et al., 2017).

Antibody dilution for immunoblotting was often 1:1000 unless specified.

Immunoprecipitation was performed as follows. HeLa Tet-On cells stably expressing Myc-Ska3 cells were lysed with lysis buffer (25 mM Tris–HCl at pH 7.5, 50 mM NaCl, 5 mM MgCl_2_, 0.1% NP-40, 1 mM DTT, 0.5 μM okadaic acid, 5 mM NaF, 0.3 mM Na_3_VO_4_ and 100 units ml^−1^ Turbo-nuclease (Accelagen). After a 1-hr incubation on ice and then a 10-min incubation at 37 °C, the lysate was cleared by centrifugation for 15 min at 4 °C at 20,817*g*. The supernatant was incubated with the antibody beads for 2 hr at 4 °C. The Myc-antibody coupled beads (Thermo Scientific, 20168) were washed four times with wash buffer (25 mM Tris–HCl at pH 7.5, 50 mM NaCl, 5 mM MgCl_2_, 0.1% NP-40, 1 mM DTT, 0.5 μM okadaic acid, 5 mM NaF, 0.3 mM Na_3_VO_4_). The proteins bound to the beads were dissolved in SDS sample buffer, resolved by SDS–PAGE and subject to mass-spectrometric analysis.

### Immunofluorescence and chromosome spread

For regular immunostaining in **Figures 4A** and **S3A**, mitotic cells were collected and then spun onto with a Shandon Cytospin centrifuge. Cells were immediately fixed with 4% ice-cold paraformaldehyde for 4 min. and then extracted with ice-cold PBS containing 0.2% Triton X-100 for 2 min. Cells were washed with PBS containing 0.1% Triton X-100 and then incubated with primary antibodies (1:1000 dilution) overnight at 4 °C. After washed with PBS containing 0.1% Triton X-100, cells were incubated at room temperature for 1hr with the appropriate secondary antibodies conjugated to fluorophores (Molecular Probes, 1:1000 dilution). After incubation, cells were washed again with PBS containing 0.1% Triton X-100, stained with 1 μg ml^−1^ DAPI and mounted with Vectashield.

For Chromosome spreads and immunostaining in **Figures 4C** and **S3D**, mitotic cells were swelled in a pre-warmed hypotonic solution containing 50 or 75 mM KCl for 15 min at 37 °C and then spun onto slides with a Shandon Cytospin centrifuge. Cells were first extracted with ice-cold PBS containing 0.2% Triton X-100 for 2 min and then fixed in 4% ice-cold paraformaldehyde for 4 min. Cells were washed with PBS and then incubated with primary antibodies (1:1000 dilution) overnight at 4 °C. Cells were washed with PBS containing 0.1% Triton X-100 and incubated at room temperature for 1h with the appropriate secondary antibodies conjugated to fluorophores (Molecular Probes, 1:1000 dilution). After incubation, cells were washed again with PBS containing 0.1% Triton X-100, stained with 1 μg ml^−1^ DAPI and mounted with Vectashield.

The images were taken by a Nikon confocal microscope with a ×60 objective. Image processing was carried out with ImageJ and Adobe Photoshop. Quantification was carried out with ImageJ.

For quantification of GFP-Ska3 on kinetochores in **Figures 4A, 4C**, **S3A**, and **S3D**, a mask was generated to mark the kinetochore on a chromosome. After background subtraction, the intensities of GFP-Ska3 and ACA signals within the mask were obtained in number. Relative intensity in (**Figures 4B, 4D**, **S3B**, and **S3D**) was derived from the intensity of GFP-Ska3 signals normalized to the one of ACA signals and plotted with the GraphPad Prizm software.

### Time-lapse microscopy

HeLa-Teton cells stably expressing H2B-mCherry were treated as indicated (**Figure 5**), and then long-term imaging was performed. Images were collected every 6 min for 12-15 hrs using a Nikon confocal microscope equipped with an environment chamber that controls temperature and CO_2_, and 20X objective. Image panels displaying the elapsed time between consecutive frames were assembled using the software designed for Nikon confocal microscope. The time taken for each cell to progress from nuclear envelope breakdown (NEB) to anaphase onset (chromatid separation) was calculated in minutes and plotted in GraphPad Prism. The experiments in (**Figure 5**) were repeated for at least two times and the results were highly reproducible. Quantification was performed based on the results from a single experiment. Average and standard deviation was calculated using GraphPad Prism.

For live-cell imaging in **Figure 4E**, GFP-Ska3 WT and 6A were expressed in HeLa Tet-On cells. After treated with MG132 for 30 min, cells were immediately subject to imaging using a Nikon confocal microscope. Images were processed in ImageJ and Adobe Photoshop.

### Protein purification

The full-length Ska3 was subcloned into pGEX-6p-1(GE Healthcare) vectors with an N-terminal GST tag with a 3C-cleavage site. The full-length Ska1-Ska2 were subcloned into the modified pET vector (Novagen) with or without 6x His tag. All Ska3 mutants were generated with standard two-step methods using PCR and Dpn1 and confirmed by DNA sequencing. For full-length Ska complex, GST-Ska3 was expressed in *E.coli* BL21 (DE3) cells, cultured in Luria-Bertani medium, Untagged or 6×His-tagged Ska1-Ska2 was cultured in Terrific Broth medium. All the cells were induced by 0.2 mM IPTG at 16°C overnight. Two sorts of cells were harvested and mixed for co-purified, and then disrupted by high-pressure homogenizer (ATX Engineering) in the PBS buffer. The cell lysates were clarified by centrifuged at 35000 g for 30 min at 4 °C. The supernatant was added GST agarose beads (GE Healthcare) and rotated at 8 °C for 1.5 hr, the protein-bound beads were then transferred into gravity columns, washed with PBS extensively, and treated with 3C protease overnight at 4 °C to remove the GST tags. After being dialyzed against 25 mM Tris pH 8.0, 50 mM NaCl and 1 mM DTT. Subsequently, the harvested Ska complex was further purified by anion exchange chromatography (HiTrap Q FF, GE Healthcare) and gel filtration chromatography (Superdex 200 10/300 GL, GE Healthcare). The purified Ska complex was finally concentrated to 15~20 mg/ml in buffer containing 20 mM Tris pH 8.0, 150 mM NaCl and 1 mM DTT, and stored at −80 °C for later use.

GST-Nuf2-Hec1 complexes, GST-Spc24-Spc25 complexes and Ndc80 Bosai, were purified as previously described (Zhang et al., 2017). To remove GST tags, cell lysates of these complexes were firstly treated with GST agarose beads (GE Healthcare). And then the protein-bound beads were transferred into gravity columns and treated with 3C protease overnight at 4 °C. The eluted complexes were subject to gel filtration chromatography (Superdex 200 10/300 GL, GE Healthcare) for further purification.

### *In vitro* binding and kinases assays

In **Figures 2B** and **2C**, 5 ug-10 ug GST fusion protein (GST-Nuf2:Hec1/GST-Spc24:Spc25) in buffer containing 20 mM Tris pH 8.0, 150 mM NaCl and 1 mM DTT, was bound to previously-equilibrated GST beads at 8 °C for 1 hr, then incubated with Ska complex or mutants proteins at 8 °C for 1.5 hr, and washed three times with buffer containing 20 mM Tris pH 8.0, 300 mM NaCl, 1 mM DTT and 0.02% TritonX-100. The proteins retained on the beads were resolved with SDS-PAGE and stained with Coomassie Blue. GST proteins were also included as control.

In **Figure 2A**, purified Ska complexes (WT and 6A) were treated with CDK1-CyclinB1 plus 10 mM ATP with or without 100 uM RO3306 in kinase buffer containing 20 mM Tris pH 8.0, 50mM NaCl, 10 mM MgCl2 and 1 mM DTT at room temperature for 1 hr. Then Ska complexes were incubated with GST beads pre-bound with the GST-Nuf2-Hec1 complex for 1 hr at 8 °C. After washed with kinase buffer containing 0.02%Trition, the proteins bound with beads were resolved by SDS-PAGE stained with Coomassie Brilliant Blue.

### QUANTIFICATION AND STATISTICAL ANALYSIS

Image J was used to obtain numeric intensities of experimental subjects under investigation. In the experiments of **Figures 4A, 4C**, **S3A** and **S3D**, six kinetochores were randomly selected from each cell. A mask was generated to mark centromeres on the basis of ACA staining in the projected image. After background subtraction, the mean intensity for objects in the mask in each channel was measured. These values were then normalized by the intensity of ACA signals and plotted with GraphPad Prism or Microsoft Excel. All the experiments were repeated at least twice or three times. Quantification was usually performed based on the results from a single experiment unless specified. Differences were assessed using ANOVA followed by pairwise comparisons using Tukey’s test. All the samples analyzed were included in quantification. Sample size was recorded in figures and their corresponding legends. No specific statistical methods were used to estimate sample size. No methods were used to determine whether the data met assumptions of the statistical approach.

